# APOL1 kidney risk variants in glomerular diseases modeled in transgenic mice

**DOI:** 10.1101/2023.03.27.534273

**Authors:** Teruhiko Yoshida, Khun Zaw Latt, Briana A. Santo, Shashi Shrivastav, Yongmei Zhao, Paride Fenaroli, Joon-Yong Chung, Stephen M. Hewitt, Vincent M. Tutino, Pinaki Sarder, Avi Z. Rosenberg, Cheryl A. Winkler, Jeffrey B. Kopp

**Affiliations:** Kidney Disease Section, Kidney Diseases Branch, NIDDK, NIH, Bethesda, MD; Department of Pathology and Anatomical Sciences, Jacobs School of Medicine & Biomedical Sciences, University at Buffalo, Buffalo, NY; Frederick National Laboratory for Cancer Research, NCI, NIH, Frederick, MD; Department of Pathology, Johns Hopkins Medical Institutions, Baltimore, MD; S.C. Nefrologia e Dialisi, AUSL-IRCCS, Reggio Emilia, Italy; Center for Cancer Research, NCI, NIH, Bethesda, MD; College of Medicine, University of Florida, Gainesville, FL

## Abstract

*APOL1* high-risk variants partially explain the high kidney disease prevalence among African ancestry individuals. Many mechanisms have been reported in cell culture models, but few have been demonstrated in mouse models. Here we characterize two models: (1) HIV-associated nephropathy (HIVAN) Tg26 mice crossed with bacterial artificial chromosome (BAC)/APOL1 transgenic mice and (2) interferon-γ administered to BAC/APOL1 mice. Both models showed exacerbated glomerular disease in APOL1-G1 compared to APOL1-G0 mice. HIVAN model glomerular bulk RNA-seq identified synergistic podocyte-damaging pathways activated by the APOL1-G1 allele and by HIV transgenes. Single-nuclear RNA-seq revealed podocyte-specific patterns of differentially-expressed genes as a function of APOL1 alleles. Eukaryotic Initiation factor-2 pathway was the most activated pathway in the interferon-γ model and the most deactivated pathway in the HIVAN model. HIVAN mouse model podocyte single-nuclear RNA-seq data showed similarity to human focal segmental glomerulosclerosis (FSGS) glomerular bulk RNA-seq data. Furthermore, single-nuclear RNA-seq data from interferon-γ mouse model podocytes (*in vivo*) showed similarity to human FSGS single-cell RNA-seq data from urine podocytes (*ex vivo*) and from human podocyte cell lines (*in vitro*) using bulk RNA-seq. These data highlight differences in the transcriptional effects of the *APOL1*-G1 risk variant in a model specific manner. Shared differentially expressed genes in podocytes in both mouse models suggest possible novel glomerular damage markers in *APOL1* variant-induced diseases. Transcription factor *Zbtb16* was downregulated in podocytes and endothelial cells in both models, possibly contributing to glucocorticoid-resistance. In summary, these findings in two mouse models suggest both shared and distinct therapeutic opportunities for APOL1 glomerulopathies.

**Significance statement:** Coding variants in APOL1, encoding apolipoprotein L1, contribute to kidney disease in individuals with African ancestry. The mechanisms for glomerular injury remain incompletely understood. We studied two transgenic mouse models, HIV-associated nephropathy and interferon-γ administration. Using glomerular and single-nuclear RNA sequencing, we identified genes differentially expressed among mice with kidney risk alleles (G1) and the common variant (G0). Both models exhibited up-regulation of genes that indicated podocyte damage with risk alleles compared to the common variant. One gene down-regulated in both models was Zbtb16, encoding a transcription factor, that may contribute to glucocorticoid-resistance. Overall, the findings suggest both shared and distinct alterations in the two disease models.

## Introduction

*APOL1* high-risk variants discovered in 2010^1, 2^ explain a substantial proportion of the high prevalence of kidney diseases in individuals with sub-Saharan African ancestry.^3^ Carriage of two *APOL1* high-risk variants account for much of the excess risk for end stage kidney disease (ESKD), focal segmental glomerulosclerosis (FSGS) and HIV-associated nephropathy (HIVAN) among African Americans, 12–14% of whom carry high-risk genotypes.^4^ The utility of *APOL1* genotype testing is under consideration in various clinical settings and APOL1 inhibition is under investigation in clinical trials.^5–7^

The molecular mechanisms by which *APOL1* high-risk variants damage glomerular cells remain incompletely understood. While many mechanisms have been reported in cell culture models, few have been demonstrated to be active in transgenic mouse models.^3, 8^ APOL1 is believed to cause podocyte damage primarily via endogenous expression, as reported in podocyte-specific over-expression mouse models.^9, 10^ There have been several reports investigating mechanism of APOL1 disease using a bacterial artificial chromosome (BAC)/APOL1 transgenic mouse model, which mimics human *APOL1* expression under the regulation of the human *APOL1* promotor and other regulatory genetic elements.^11–16^

Of the many APOL1-associated kidney diseases, HIVAN has the strongest association with *APOL1* high-risk genotypes, with odds ratios of 29 and 89 in the USA and South Africa, respectively.^17, 18^ Here we report molecular pathways using perhaps the most relevant translational mouse model of APOL1 expression (BAC/APOL1) crossed with HIVAN (Tg26) mouse model. We also induced acute glomerular injury by administrating exogenous interferon-γ to BAC/APOL1 mice to drive increased APOL1 expression. We identify shared and discrete APOL1 high risk-mediated pathways opening up the potential for generic and disease specific therapeutic interventions.

## Methods

### Mice

Mouse experiments were conducted in accordance with the National Institutes of Health Guide for the Care and Use of Laboratory Animals and were approved in advance by the NIDDK Animal Care and Use Committee (Animal study proposal, K097-KDB-17 & K096-KDB-20). We studied both male and female mice. Mice were housed in cages on a constant 12-hour light/dark cycle, with controlled temperature and humidity and food and water provided ad libitum. Sample sizes for experiments were determined without formal power calculations. Mice with large cutaneous papilloma were excluded from the study. Investigators were not masked to group allocation but were masked when assessing outcome.

We used human *APOL1* gene locus transgenic mice (BAC/APOL1 mice)^13, 14^, generated using a bacterial artificial chromosome containing APOL1-G0 or variant G1 or G2 alleles. As previously described, ∼47 kb human DNA, encompassing only the human APOL1 gene with 5ʹ and 3ʹ flanking regions (including exons 1 and 2 of APOL2 and 3ʹ region including exons 39–41 of part of MYH9 gene), was isolated and subcloned from human BAC clone (ENST00000397278, which corresponds to NM_003661). Individual G0, G1, and G2 BAC subclones were injected into 129SvJ/ B6N F1 mouse embryos and the founder mice were subsequently backcrossed into the 129SvJ background.

For the HIV-associated nephropathy (HIVAN) model mouse, we bred hemizygous BAC/APOL1-G0/G1/G2 mice with Tg26 mice to obtain F1 generation of BAC/APOL1xTg26 dual transgenic mice for experiments. We measured weights and collected random urine at 6 and 9 weeks of age. We used 9-week-old mice for phenotype characterizations, including glomerular extraction and collection of kidneys and blood as described below.

For interferon-γ model experiments, we injected a single dose of recombinant murine interferon-γ (CYT-358, ProSpec, East Brunswick NJ) in phosphate-buffered saline (PBS) at 1.125 × 10^7^ U/kg body weight, retro-orbitally under isoflurane anesthesia (0 hour).^16^ Urine was collected every 24 hours using metabolic cages, beginning 24 hours prior to interferon injection and ending 72 hours after injection. Selected mice were euthanized at 24hours for further experiments.

For triple intervention experiments, we administered murine interferon-γ, puromycin aminonucleoside, and basic fibroblast growth factor^14^ Mice received injections on days −1 and +1 with interferon-γ at a dose of 10^6^ U/kg body weight. On day 0, mice received puromycin aminonucleoside (Millipore-Sigma, Burlington, MA) by intraperitoneal administration, at a dose of 300 mg/kg body weight. On days 0 and 2, mice received basic fibroblast growth factor, 5 µg intravenously (Kaken Pharmaceutical, Tokyo, Japan). Glomerular filtration rate was measured at day 10, as described below. Urine was collected at day −2, day 7, and day 14 for 24 h using metabolic cages. Mice were euthanized for further experiments on day 14.

### Measurement of Mouse Glomerular Filtration Rate (**GFR**)

GFR was measured ten days after puromycin aminonucleoside injection of triple intervention model. After anesthesia with 2.5% isoflurane, the dorsal hair was shaved. A fluorescence sensor and a battery pack (NIC-Kidney, Mannheim Pharma & Diagnostics, Mannheim, Germany) was placed on the dorsum and secured with surgical tape. FITC-sinistrin (Medi Beacon, St. Louis, MO) was dissolved in 100 µl PBS (15 mg/100g body weight) and injected intravenously or retro-orbitally. Fluorescent signals were measured for approximately one hour, after which the sensor and battery were removed for analysis. Data were analyzed according to the manufacturer’s instructions. Briefly, GFR was calculated using the half-life (t_1/2_) derived from the rate constant of the single exponential elimination phase of the fluorescence-time curve and a semi-empirical mouse-specific conversion factor established previously.^19^

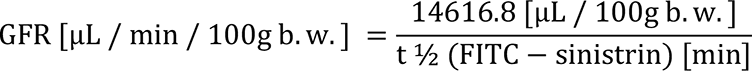

### Urinary Albumin and Creatinine Measurement

We determined the urinary albumin levels with Albuwell M (Mouse Albumin ELISA) (1011, Ethos Biosciences, Logan Township, NJ). We measured the urine creatinine concentration with Creatinine Companion kit (1012, Ethos Biosciences). Albuminuria was determined as the ratio of urinary albumin to creatinine. All procedures were performed in accordance with the manufacturers’ protocols.

### In situ Hybridization

Mouse kidney tissues were fixed with 10% buffered formalin for 24 hours and embedded in paraffin. Chromogenic in situ detection was performed on tissue sections from the mouse formalin-fixed paraffin-embedded (FFPE) blocks using the RNAscope in situ hybridization (Advanced Cell Diagnostics, Biotechne, Minneapolis, MN). Briefly, 5 μm FFPE tissue sections were de-paraffinized, boiled with RNAscope Target Retrieval Reagent for 15 min at 99°C, and protease digested at 40°C for 30 min. This was followed by hybridization for 2 h at 40°C with probe-Hs-*APOL1*-O1 (#439871, Advanced Cell Diagnostics). In addition, Probe-Mm-PPIB (#313911) and Probe-DapB (#310043) were used for positive and negative control, respectively. Probes were detected with RNAscope 2.5 HD Reagent Kit (Brown) (#322310).

Fluorescent in situ detection of mRNA was performed using RNA probe-Hs-APOL1-No-XMm (#459791), Mm-Nphs1 (#433571), Nell2 (#1093741), St6galnac3 (#1170041), Ccn2 (#1170051). Specific probe binding sites were visualized using RNAscope Hiplex12 Reagents Kit (488, 550, 650) v2 (Advanced Cell Diagnostics, Biotechne) (#324419). A Nikon Sora microscope in the NIDDK Advanced Light Microscopy Imaging Analysis Core facility was used for fluorescence imaging.

### Immunohistochemistry

FFPE tissue sections were deparaffinized and rehydrated. Antigen retrieval was performed by heating in citrate-buffered medium for 15 min in a 99°C hot water bath. Tissues were blocked by 2.5% normal horse serum. Sections were incubated for one hour at room temperature with 5 μg/ml of primary antibody for APOL1 (5.17D12, rabbit monoclonal) that was kindly provided by Genentech (South San Francisco, CA). ^20^ Sections were processed following ImmPRESS HRP Universal Antibody (horse anti-mouse/rabbit IgG) Polymer Detection Kit and ImmPACT DAB EqV Peroxidase (HRP) Substrate (Vector Laboratories, Burlingame, CA) protocol, and counter-stained with hematoxylin.

### Immunofluorescence Staining

Mouse kidney tissues were harvested and embedded in OCT-compound and frozen. Frozen blocks were sectioned at 5 μm. Sections were hydrated in PBS and blocked with 10% bovine serum albumin (BSA) / PBS for one hour in a humidified chamber. Alexa Fluor 647 anti-mouse F4/80 Antibody (123121, BioLegend, San Diego, CA, 1:25 dilution) were applied and incubated overnight at 4°C. After DAPI staining and mounting, slides were imaged by fluorescence microscopy.

### Confocal Microscopy

A Yokogawa CSU-W1 SoRa spinning disk confocal scanhead (Yokogawa, Sugar Land, TX), with 50-micron pinhole (standard confocal mode, no SoRa), mounted on a Nikon Ti2 microscope running NIS-elements 5.21.02 software (Nikon Instruments, Melville, NY), was used to collect tiles of multi-color fluorescence images. Fluorescence image channels were obtained sequentially, while sharing the Yokogawa T405/488/568/647 dichroic. For DAPI fluorescence: excited by the 405nm laser, emission was filtered by ET455/58 (Chroma, Technology Corp, Bellows Falls, VT). For green fluorescence: excited by the 488nm laser, emission filtered by ET520/40 (Chroma). For orange fluorescence: excited by the 561nm laser; emission filtered by ET605/52 (Chroma). For far red fluorescence: excited by the 640nm laser; emission filtered by ET655LP (Chroma). Images used for quantitation of stain prevalence were acquired with the Nikon Apo TIRF 60x/1.49 Oil DIC N2 objective lens, producing a confocal section thickness of 1.2 microns for all fluorescence channels.

### Transmission Electron Microscopy and Foot Process Effacement Evaluation

Mouse kidney sections, ∼1 mm^3^, were fixed for 48 h at 4°C in 2.5% glutaraldehyde and 1% paraformaldehyde in 0.1M cacodylate buffer (pH 7.4) and washed with cacodylate buffer three times. Tissues were fixed with 1% OsO_4_ for two hours, washed again with 0.1M cacodylate buffer three times, washed with water and placed in 1% uranyl acetate for one hour. Tissues were serially dehydrated in ethanol and propylene oxide and embedded in EMBed 812 resin (Electron Microscopy Sciences, Hatfield, PA). Thin sections (∼80 nm) were obtained with a Leica Ultracut-UCT ultramicrotome (Leica, Deerfield, IL), placed onto 300 mesh copper grids and stained with saturated uranyl acetate in 50% methanol, followed by lead citrate. Grids were viewed with a JEM-1200EXII electron microscope (JEOL Ltd, Tokyo, Japan) at 80 kV and images were recorded on the XR611M, mid-mounted, 10.5 M pixel, CCD camera (Advanced Microscopy Techniques Corp, Danvers, MA). We evaluated for foot processes effacement the filtering surface of open capillary loops, using image processing software (ImageJ, Viewpoint Light). We measured the length of the outer surface of the glomerular basal membrane. Overlying foot processes were counted by hand. A ratio of foot process number to glomerular basement membrane length (GBM) length was computed for each picture.

### Mouse Kidney Pathological Evaluation

FFPE mouse kidney tissue sections, cut at 4 µm, were stained with hematoxylin and eosin, periodic acid-Schiff (PAS) reagents for routine and quantitative histological assessment.

### Estimation of Glomerular Podocyte Count

‘PodoCount’^21^, a computational tool for whole slide podocyte estimation from digitized histologic sections, was used to detect, enumerate, and characterize podocyte nuclear profiles in the glomeruli of immunohistochemically labeled mouse kidney sections. FFPE tissues (2 µm thickness) were immuno-stained for p57^kip2^, a marker of podocyte terminal differentiation (ab75974, Abcam, Cambridge, UK), and detected with horse radish peroxidase (RU-HRP1000, Diagnostic BioSystems, Pleasanton, CA) and diaminobenzidine chromogen substrate (BSB0018A, Bio SB, Santa Barbara, CA). PAS post-stain was applied without hematoxylin counterstain. PodoCount uses a combination of structural segmentation steps that employ thresholding, a convolutional neural network^22^, and stain deconvolution; morphological image processing approaches to refine segmentation; and literature-informed feature engineering steps to compute various histologic podometrics^23^ with correction for section thickness.^24^ In this study, PodoCount was used to assess mean glomerular podocyte count and podocyte density per mouse.

### Glomerular Isolation

Mice were anesthetized with 2, 2, 2-trimbromoethanol (Avertin), the abdominal aorta was exposed and, a cannula was inserted under microscopy into the aorta with a PE-10 tube attached to a PE-50 tube. After clipping the celiac trunk, superior mesenteric artery and aorta proximal to renal arteries, kidneys were perfused with PBS via the cannula. A small incision was made in the proximal left renal vein to assure good reagent perfusion and the left kidney vessels were clipped. The right kidney was perfused twice with PBS, containing 10ul of Dynabeads M-450 tosylactivated (14013, Invitrogen, Waltham, MA). The right kidney was extracted for glomerular isolation and left kidney was extracted for pathological characterization.

The right kidney was placed in Hanks’ Balanced Salt solution (HBSS) on ice. The kidney was minced with razor blades. Kidney fragments were placed in a solution containing 4mg/ml of collagenase A (10103586001, Roche, Mannheim, Germany) and 200 units/ml of DNase I (04716728001, Roche) for 30 mins in 37°C with 1500rpm agitation. A 100-µm strainer (542000, greiner bio-one, Frickenhausen, Germany) and Dynal (Thermofisher, Waltham, MA) magnet was used to isolate glomeruli washed three times with PBS. For bulk-RNA-seq samples, 500 µl QIAzol (QIAGEN, Hilden, Germany) was added to 1.5ml tube containing glomeruli from 1/6 of each kidney. For single-nuclear RNA-seq samples, purified glomeruli were snap frozen in 1.5ml tubes.

### RT-qPCR

Total RNA was extracted using RNeasy Plus Universal Kit (73404, QIAGEN) following the manufacturer’s protocol, including removal of genomic DNA. RNA (1 µg) was reverse transcribed using GoScript Reverse Transcription System (A5001, Promega, Madison, WI). cDNA was measured by quantitative PCR with FastStart Universal SYBR Green Master (Rox) (04913850001, Sigma-Aldrich, St. Louis, MO) using QuantStudio 6 (ThermoFisher). Relative mRNA levels were quantified by the ddCT method, using β-actin as an endogenous control. Primers used are summarized in **Supplemental Table S1**.

### Bulk Glomerular RNA-Sequencing

Isolated glomerular tissues from 8 line (WT, G0, G1, G2, Tg26, G0xTg26, G1xTg26, G2xTg26; n=5, 3, 4, 3, 5, 5, 5, 4, respectively) were homogenized in QIAzol. Total RNA was extracted using RNeasy Plus Universal Kit (73404, QIAGEN) following the manufacturer’s protocol, including removal of genomic DNA. RNA samples were prepared sequencing at the sequencing facility, Frederick National Laboratory for Cancer Research, NCI, Frederick, MD.

Samples were pooled and sequenced on NovaSeq6000 S1 flow cell using Illumina TruSeq Stranded mRNA Library Prep and paired-end sequencing with read length 101bps (2x101 cycles). Samples had 47 to 67 million reads passing a quality filter and more than 92% of bases calls had a quality score above Q30. Sample reads were trimmed to remove adapters and low-quality bases using Cutadapt v1.18. Trimmed reads were mapped to the reference genome (Mouse mm10) and to transcripts (Ensembl v96 annotation) using STAR aligner v2.7.0f. Gene expression quantification was performed using RSEM v1.3.1 tool. The sequencing and mapping statistics of bulk glomerular RNA-seq are in **Supplemental Table S2**.

### Bulk Glomerular RNA-sequencing Gene Enrichment and Network Analysis

DESeq2^25^ was used for differential expression analysis from raw count data and normalized data were used for gene set enrichment analysis. GSEA v4.1.0^26, 27^ was used for pathway enrichment analysis. Pathway analysis, including canonical pathway analysis and upstream regulator analysis, was performed using the QIAGEN Ingenuity Pathway Analysis (IPA) software. The WGCNA package^28^ was used for the network analysis with the normalized counts data from APOL1-G0, G1, G0xTg26, G1xTg26 samples by DESeq2, to identify modules related to *APOL1* genotype and the Tg26 transgene. Network building and visualization of genes in the saddlebrown module were conducted by NetworkAnalyst 3.0 (https://networkanalyst.ca/).^29^ Intersection of lists were mapped on STRING Interactome (900 Confidence score with experimental evidence).

### Single-nucleus RNA-sequencing

Nuclei from isolated five glomeruli sample types (WT, G0xTg26, G1xTg26, G0+interferon-γ, G1+interferon-γ) were prepared as follows^30^. Snap frozen isolated glomeruli were lysed in EZlysis buffer (#NUC101-1KT, Sigma, Darmstadt, Germany) and homogenized 30 times using a loose Dounce homogenizer and 5 times in a tight pestle. After 5 min incubation, the homogenate was passed through a 40 µm filter (43-50040, PluriSelect, El Cajon, CA) and centrifuged at 500g at 4°C for 5 min. The pellet was washed with EZlysis buffer and again centrifuged at 500g at 4°C for 5 min. The pellet was resuspended with DPBS with 1% FBS and passed through a 5 µm filter (43-50005, PluriSelect) to make final nuclei prep for loading on to a 10x Chromium Chip G (10X Genetics, Pleasanton, CA) for formation of gel beads in emulsion (GEM).

Single nuclear isolation, RNA capture, cDNA preparation, and library preparation were performed following the manufacturer’s protocol (Chromium Next GEM Single Cell 3’ Reagent Kit, v3.1 chemistry, 10x Genomics). Prepared cDNA libraries were sequenced at Frederick National Laboratory for Cancer Research. All samples had sequencing yields of at least 183 million reads with over 95.7% of bases in the barcode regions had Q30 or above and at least 93.5% of bases in the RNA read had Q30 or above. More than 95.5% of bases in the unique molecular identifiers (UMI) had quality scores Q30 or above. Analyses were performed with the Cell Ranger v6.1.2 software (10x Genomics) using the default parameters with pre-mRNA analysis turned on. The reference was built from mouse (mm10) reference genome, supplemented with HIV-1 viral sequences and human *APOL1* sequences. The sequencing and mapping statistics of single-nucleus RNA-seq are in **Supplemental Table S3**.

### Single-nucleus RNA-sequencing Analysis

Removal of ambient RNA was conducted by SoupX (version 1.5.2)^31^ following the default protocol by “autoEstCont” and “adjustCounts” functions. After removal of ambient RNA, integration of single-nucleus gene expression data was performed using Seurat (version 4.0.5) and SeuratData (version 0.2.2)^32^ after filtering out nuclei with the any of the following features: detected genes numbers <200 or >4000, total RNA count >15,000, or mitochondrial transcript percentage >20%. After filtering, 20,276 nuclei remained for downstream analysis.

Clustering of the combined data used the first 30 principal components at a resolution of 0.6 and identified 21 distinct cell clusters. Cell types were identified by expression levels of known marker genes. After clusters from tubules, fibroblasts and doublets were removed, there remained 16,030 nuclei in 12 clusters, and these were analyzed further (**Supplemental Figure 3**). Differential expression analysis was performed using the “Findmarkers” function in Seurat, using default parameters. DEG were identified for in each paired comparison (HIVAN, G1xTg26 vs G0xTg26; interferon-γ, G1+interferon-γ vs G0+interferon- γ), using a cut-off of adjusted P <0.05. Pathway analysis was performed using QIAGEN Ingenuity Pathway Analysis (IPA)^33^ software, using DEG sets as inputs. Cell-cell interaction analysis was performed using CellChat (version 1.5.0)^34^ with mouse protein-protein interactions loaded.

### Gene set enrichment analysis of human glomerular RNA-seq data with APOL1 genotype using custom gene sets from DEG sets

Normalized human glomerular RNA-seq data were obtained from the website “APOL1 Portal” (http://APOL1portal.org).^35^ Original data were from Nephrotic Syndrome Study Network (NEPTUNE)^36^, a biopsy-based cohort of patients with proteinuric kidney diseases.

Subject inclusion criteria for the study included a histologic diagnosis of FSGS, available glomerular RNA-seq data, known *APOL1* genotype, and self-identified “Black/ African American” race or genotype-predicted African continental ancestry.^35^ DEG sets were obtained from single-nucleus RNA-seq analysis and were converted to custom human gene lists. GSEA analysis was conducted by GSEA v4.1.0, comparing data from *APOL1* high-risk subject (defined as those carrying two *APOL1* kidney risk alleles) (n=16) and data from *APOL1* low-risk subject (defined as those carrying zero or one *APOL1* kidney risk alleles) (n=14).

### Pathway analysis to compare transcriptome of APOL1-G1/G2 podocytes and APOL1-G0/G0 podocytes

Single-cell RNA-seq of urinary podocytes from FSGS subjects^37^ and bulk RNA-seq of podocytes *in vitro* (human urine-derived podocyte-like cell lines, HUPEC)^38^ were reported previously. We reanalyzed those data to compare pathways between APOL1-G1/G2 podocytes and APOL1-G0/G0 podocytes. The original data were obtained from GSE176465^37^ and GSE194337.^38^ DEGs were identified for each paired comparison (APOL1-G1/G2 vs APOL1-G0/G0). DEG sets as input for IPA analysis were defined using cut-offs as follows: adjusted P <0.05 for differentiated HUPECs, and P < 0.05 without multiple testing correction for urine podocytes from FSGS subjects. Pathway analysis was performed using QIAGEN Ingenuity Pathway Analysis (IPA)^33^ software, using the three DEG sets as inputs.

## Results

### BAC/APOL1-G1xTg26 (HIVAN) mice had accelerated glomerular injury

We established a dual transgenic mouse model BAC/APOL1xTg26 to elucidate the effect of *APOL1* genotype in the context of HIV-associated nephropathy (HIVAN). By histomorphology, we observed augmented glomerular damage in BAC/APOL1-G1xTg26 mice compared with other genotypes (**Figure 1A-H**). We confirmed APOL1 expression by immunohistochemistry (**Supplemental Figure 1A-H**) and *in situ* hybridization (ISH) (**Supplemental Figure 1I-P**), which showed APOL1 protein/mRNA expression primarily in podocytes and renal endothelial cells.

**Figure 1.**
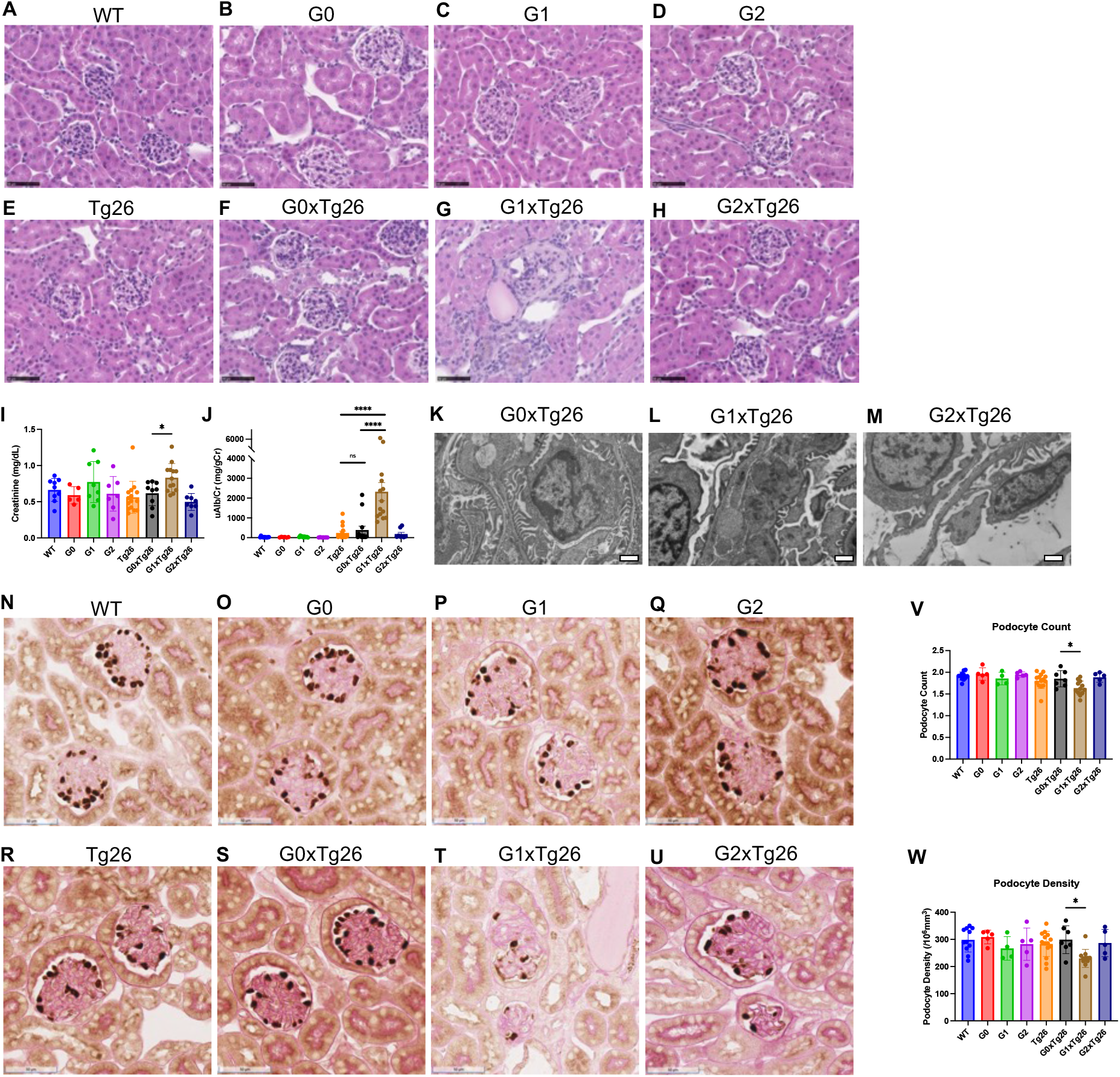
Characterization of BAC/APOL1xTg26 dual transgenic HIVAN model mouse. (**A-H**) Representative hematoxylin and eosin staining images of WT, G0, G1, G2, Tg26, G0xTg26, G1xTg26, G2xTg26 mouse kidney, noting glomerulosclerosis and microcystic tubular dilatation in G1xTg26 mice (**I**, **J**) Plasma creatinine (mg/dL), urinary albumin-to-creatinine ratio (mg/g creatinine), elevated in G1xTg26 mice compared to other mice (**K-M**) Representative electron micrographs of G0xTg26, G1xTg26, G2xTg26 kidneys, demonstrating partial foot process effacement in G1xTg26 and G2xTg26 mice (Scale bars are 100nm) (**N-U**) p57 staining of kidney showing podocyte loss and dedifferentiation in G1xTg26 mice (Scale bars are 50 μm) (**V**, **W**) Podocyte counts and densities by ‘PodoCount’ analysis showing podocyte depletion in G1xTg26 mice (one-way ANOVA; *, P<0.05; ***, P<0.001)

Plasma creatinine (**Figure 1I**) and albuminuria (**Figure 1J**) levels indicated that G1xTg26 mice had the most renal parenchymal injury, the latter suggesting primary glomerular/podocyte damage. Electron microscopy showed more prominent podocyte foot process effacement in G1xTg26 and G2xTg26 mice compared with G0xTg26 mice (**Figure 1K-M, Supplemental Figure 1Q**). Podocyte depletion in G1xTg26 mice was identified by p57 staining and podocyte estimation using ‘PodoCount’-derived podometric analysis based on podocyte count and podocyte density (**Figure 1N-W**). These data highlight an enhancement of glomerular injury, in particular podocyte injury, in a HIVAN model in an *APOL1* high-risk allele dependent manner.

### Downregulation of ribosomal genes in glomeruli from APOL1-G1 and HIVAN mice

To elucidate the molecular pathways that were dysregulated in glomeruli, we performed bulk RNA-seq from isolated glomeruli. When comparing Tg26 mice with WT mice, we found six upregulated and five down-regulated KEGG pathways (**Figure 2A, Supplemental Figure 2A, Supplemental Table S3**). The extracellular matrix-related pathway and focal adhesion pathways were enriched in Tg26 mice, and the ribosome pathway was downregulated in Tg26 mice (**Figure 2B, 2C**).

**Figure 2.**
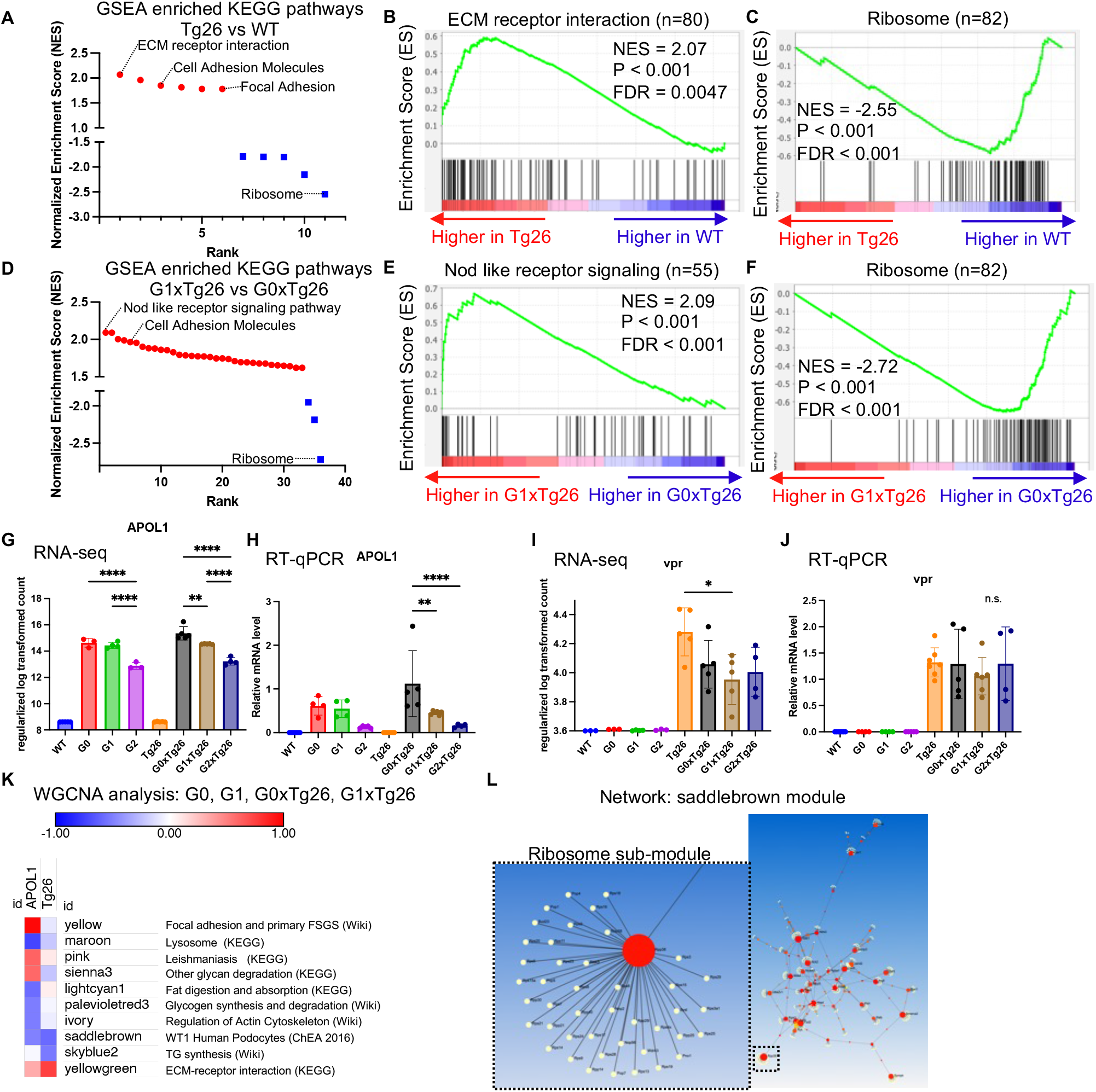
Bulk RNA-seq of glomeruli from HIVAN model mouse. (**A**) Waterfall plot showing gene set enrichment analysis results of enriched KEGG pathways comparing Tg26 and WT mice (**B**) Enrichment plot of ECM receptor interaction pathways based on bulk RNA-seq comparing Tg26 and WT mice (**C**) Enrichment plot of ribosome pathway based on bulk RNA-seq comparing Tg26 and WT mice (**D**) Waterfall plot showing gene set enrichment analysis results of enriched KEGG pathways comparing G1xTg26 and G0xTg26 mice (**E**) Enrichment plot of Nod-like receptor signaling pathway based on bulk RNA-seq comparing G1xTg26 and G0xTg26 mice (**F**) Enrichment plot of ribosome pathway based on bulk RNA-seq comparing G1xTg26 and G0xTg26 mice (**G**) Regularized log transformed counts of *APOL1* by bulk RNA-seq (**H**) Relative mRNA levels of *APOL1* by RT-qPCR (**I**) Regularized log transformed counts of *vpr* by bulk RNA-seq (**J**) Relative mRNA levels of *vpr* by RT-qPCR (**K**) Heatmap showing interactions of APOL1 and Tg26 with significant modules and representative pathway enriched in each module. Red denotes up-regulation, blue denotes down-regulation. (**L**) Network visualization of the ‘saddlebrown’ module genes layered onto the STRING interactome. The image of the ribosome sub-module is magnified, showing the complexity of this network.

When comparing G1xTg26 with G0xTg26 mice, we found 33 upregulated and three downregulated KEGG pathways (**Figure 2D, Supplemental Table S4**). The Nod-like receptor signaling pathway was the top enriched pathway in G1xTg26 mice and ribosomal pathway was again negatively enriched in G1xTg26 mice (**Figure 2E, 2F**). *APOL1* mRNA expression level in glomeruli was measured by RNA-seq and was confirmed by RT-qPCR. G0 had higher expression compared with G1, and G2 had the lowest (**Figure 2G, 2H**).

The presence of the Tg26 transgene did not increase *APOL1* expression levels, despite the greater albuminuria in G1xTg26 mice compared to G1 mice. This suggests that glomerular *APOL1* mRNA expression level was not the major driver of kidney phenotype in this BAC/APOL1xTg26 mouse model, although we cannot exclude that increases in APOL1 protein expression may have contributed. Tg26 transgene RNA expression levels were not increased by the APOL1 high-risk variant (**Supplemental Figure 2B**), as confirmed by RT-qPCR for Vpr expression level (**Figure 2I, 2J**).

### Network analysis reveals two-hit transcriptomic changes by APOL1 genotype and Tg26

WGCNA analysis was applied to the APOL1-G0, G1, G0xTg26, G1xTg26 datasets to identify modules of genes affected by *APOL1* genotype (G1 vs G0), with or without Tg26. Out of 89 modules identified, eight modules were dysregulated by APOL1-G1 compared to APOL-G0 and three modules were dysregulated by Tg26 (**Figure 2K**). The ‘saddlebrown’ module was the only module dysregulated significantly by *both APOL1* genotype and Tg26, and it was enriched genes reported by WT1 CHIP-Seq by previous report.^39^

These findings indicates that both APOL1-G1 and Tg26 transgene contribute to WT1 deactivation, suggesting podocyte dedifferentiation in these two settings. Primary FSGS and lysosome pathways genes were enriched in APOL1 related modules. Extracellular matrix pathway genes were enriched in Tg26 related modules. Network visualization of ‘saddlebrown’ module genes onto the STRING interactome showed sub-module related to ribosome KEGG pathway, which confirms ribosomal gene expression dysregulation by APOL1-G1 and Tg26 (**Figure 2L**).

### Interferon-γ induced proteinuria in BAC/APOL1 mice models acute effects of APOL1

We studied chronic and acute APOL1-induced kidney disease model on BAC/APOL1. BAC/APOL1-G0 and -G1 mice had no apparent kidney phenotype in the absence of intervention.^13^ The chronic APOL1-induced kidney disease model in these mice involved triple intervention, including interferon*-*γ, basic fibroblast growth factor, and puromycin aminonucleoside.^14^ We demonstrated glomerulosclerosis development (**Figure 3A-C**), nephrotic albuminuria (**Figure 3D, E**), reduced GFR (**Figure 3F**) and podocyte depletion (**Figure 3G, H**) after triple intervention in G1 and G2, but not in G0 transgenic mice.

**Figure 3.**
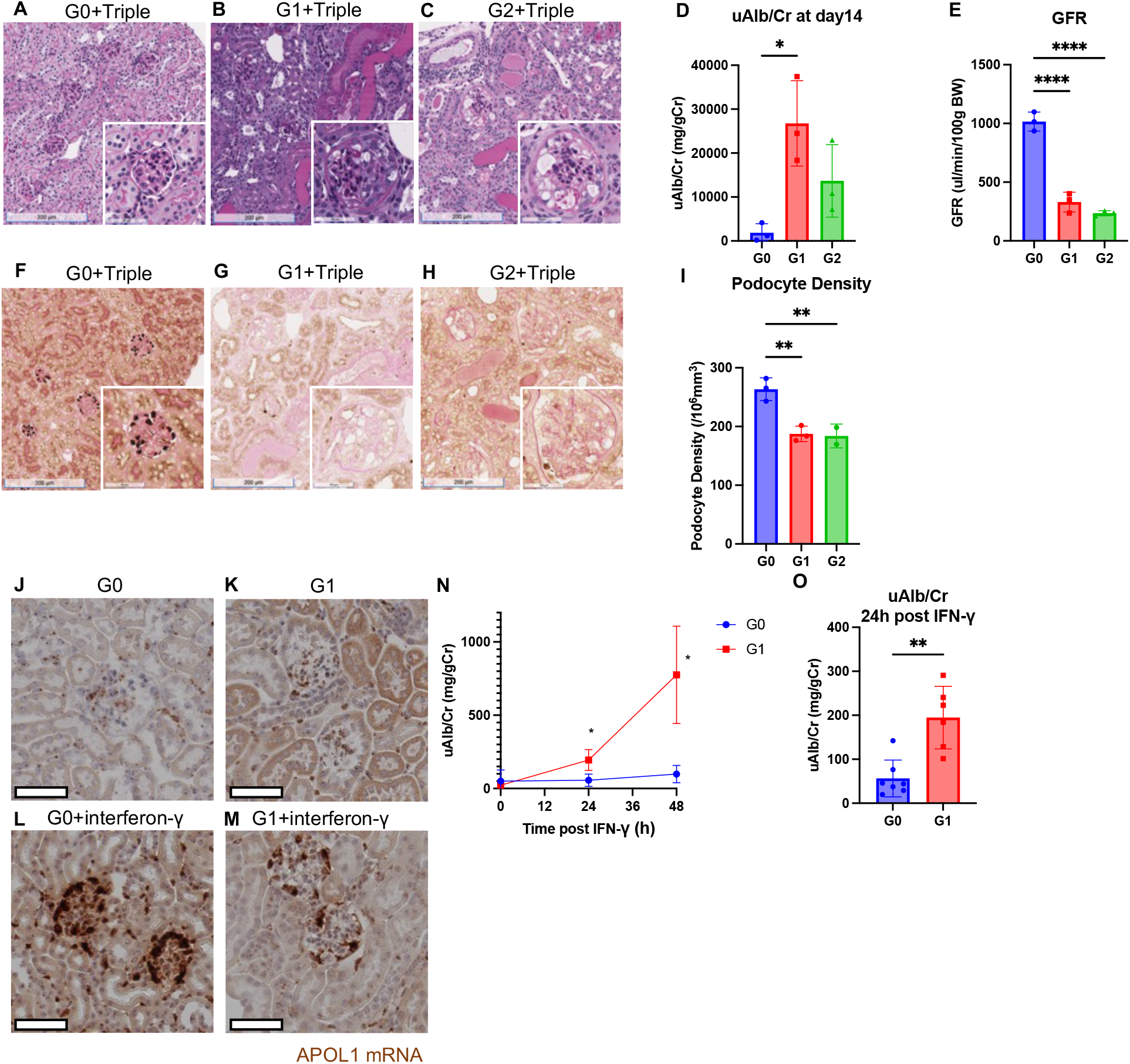
Characterization of triple intervention and interferon-γ model. (**A-C**) Representative hematoxylin and eosin staining images of G0, G1, G2 kidney underwent triple intervention (14 days post) (**D**) Urinary albumin-to-creatinine ratio at 14 days post triple intervention (**E**) Glomerular filtration ratio (GFR) measured at 10 days post triple intervention (**F-G**) Representative p57 podocyte staining images of G0, G1, G2 kidney underwent triple intervention (14 days post) (**I**) Podocyte densities by ‘PodoCount’ analysis showing podocyte loss in G1 and G2 mice (**J-M**) Representative images of ISH of *APOL1* showing enhanced expression of APOL1 post 24 hours of interferon-γ injection (**N**) Time-course measurements of urinary albumin-to-creatinine ratio (mg/g creatinine) in interferon-γ injection experiments (**O**) Urinary albumin-to-creatinine ratio at 24 hour following interferon-γ injection (one-way ANOVA; *, P<0.05; **, P<0.01; ***, P<0.001; ****, P<0.0001)

We next investigated the acute effect of APOL1-G1 on glomerular cells, using the interferon*-*γ IV model.^16^ We have shown that APOL1 expression was induced in both G0 and G0 mice (**Figure 3I-L**) but that albuminuria was induced only in G1 mice, not in G0 mice as reported (**Figure 3M**).^16^ Using metabolic cages for urine collection, we found that compared to G0 mice, G1 mice had significant albuminuria increases within the first 24 hours after administration of interferon-γ, timepoint that was earlier than in previous reports (**Figure 3N**).^16^ Therefore, we collected and characterized glomeruli from mice 24 hours after interferon*-*γ injection to investigate the effect of APOL1-G1 compared to G0 in the acute phase induced by interferon*-*γ.

### Single-nuclear RNA-seq showed podocyte damage patterns by APOL1-G1 depending on the model

To characterize molecular signatures at single cell resolution, we conducted single-nuclear RNA-seq of isolated glomeruli from WT, G0xTg26, G1xTg26, G0+interferon-γ, G1+interferon-γ mice kidney. We profiled 20276 nuclei in total. We identified 21 cell clusters shown by UMAP (**Supplemental Figure 3A, 3B**). We took subset data of the above cell clusters to focus on glomerular characterization. We annotated cell types into 12 clusters according to prior knowledge of gene expression, including three clusters of podocytes, three clusters of mesangial cells, three clusters of endothelial cells, one cluster of parietal epithelial cells (PEC), one cluster of immune cells, and one juxtaglomerular (JG) cell cluster (**Figure 4A, B**).

**Figure 4.**
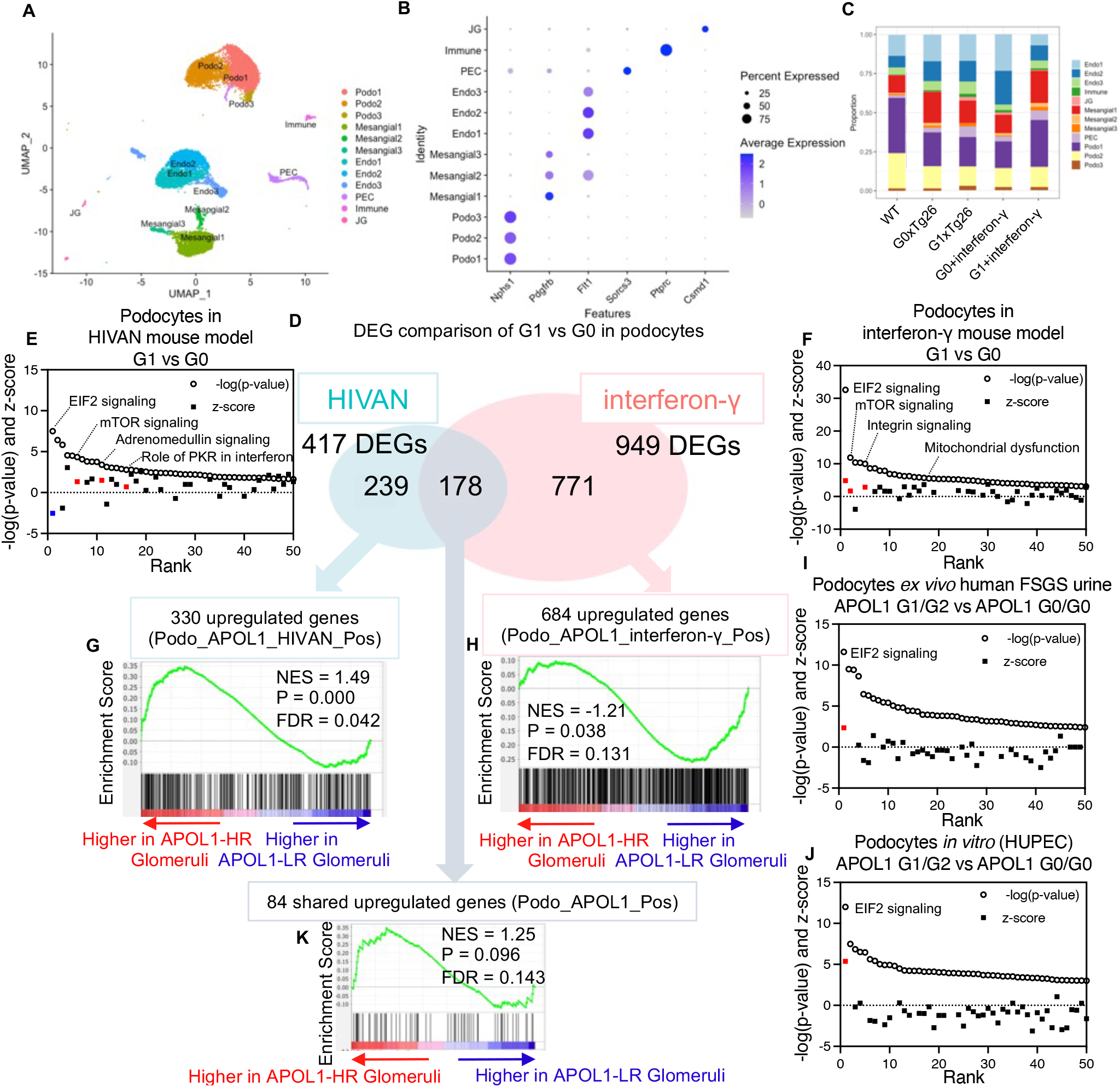
Overview of single-nucleus RNA-seq experiments. (**A**) UMAP plot of single-nuclear RNA-seq data for analysis from five samples, showing 12 clusters. (**B**) Dot plot of marker genes characteristic for each cluster (**C**) Ratio of nuclei grouped to each cluster by each sample (**D**) Venn diagram of DEGs, comparing APOL1 genotype (G1 vs G0) in both HIVAN and interferon-γ model podocytes (**E**) Waterfall plot of IPA pathways enriched by DEGs from podocytes comparing G1 and G0 in the HIVAN model (**E**) Waterfall plot of IPA pathways enriched by DEGs from podocytes comparing G1 and G0 in interferon-γ model (**G**) Enrichment plot of Podo_APOL1_HIVAN_Pos genes on human glomerular RNA-seq comparing APOL1-HR and LR (**H**) Enrichment plot of Podo_APOL1_interferon-γ_Pos genes on human glomerular RNA-seq comparing APOL1-HR and LR (**I**) Waterfall plot of IPA pathways enriched by DEGs from podocytes *ex vivo* human FSGS urine single-cell RNA-seq data comparing APOL1-G1/G2 samples and G0/G0 samples (GSE176465) (**J**) Waterfall plot of IPA pathways enriched by DEGs from podocytes *in vitro* (HUPEC) comparing APOL1-G1/G2 and G0/G0 (GSE194337) (**K**) Enrichment plot of Podo_APOL1_ Pos genes on human glomerular RNA-seq comparing APOL1-HR and LR

*APOL1* was expressed in podocytes and endothelial cells (**Supplemental Figure 3C**). HIV genes were relatively highly expressed in podocytes compared with other glomerular cell types (**Supplemental Figure 3D**). Immune cells were more frequent in G1xTg26 mice compared with G0xTg26 mice, which was confirmed by F4/80-positive macrophage staining (**Figure 4C, Supplemental Figure 3E-H**). Regarding all nine glomerular cell clusters, we regrouped these into three new cell clusters, comprised podocytes, endothelial cells and mesangial cells. Using these podocytes and endothelial cells clusters, we conducted differential expression analysis to define DEGs (adjusted p-value<0.05) comparing G1xTg26 with G0xTg26 (HIVAN) mice and G1+interferon-γ with G0+interferon-γ (interferon-γ) mice. Thus, we created four DEG sets: Podo_APOL1_HIVAN (n=417); Podo_APOL1_interferon-γ (n=949); Endo_APOL1_HIVAN (n=166); Endo_APOL1_interferon-γ (n=804) (**Figure 4D**).

When we conducted IPA analysis based on these DEG sets, we identified pathways dysregulated in each cell cluster in two different models, HIVAN and interferon-γ (**Figure 4E, F**). The EIF2 pathway was the top canonical pathway dysregulated in podocytes in both models, with different directionality. In this pathway, the interferon-γ model manifested *activation* (active Z-score 4.85), while the HIVAN model manifested *deactivation* (active Z-score -2.53). These findings suggest that the effect of APOL1 variant on the EIF2 pathway may be context dependent. When we compared DEG sets of podocytes in the two models (**Figure 4D**), we identified 178 shared DEGs in both models. Out of 178 DEGs shared, 84 DEGs were upregulated in both models (Podo_APOL1_Pos, (n=84)). There were 330 upregulated DEGs in the HIVAN model (Podo_APOL1_ HIVAN_Pos (n=330)) and 684 in the interferon-γ model (Podo_APOL1_ IFN-γ_Pos DEG gene set (n=684)). These upregulated DEGs of podocytes had positive enrichment in bulk RNA-seq data, comparing G1xTg26 with G0xTg26 mice and G1 with G0 mice (**Supplemental Figure 4A-C**).

### Comparing podocyte damage patterns by APOL1-G1 in mouse models with human data

To extrapolate the relevance of these data to human disease, we conducted gene set enrichment analysis using publicly available human glomerular RNA-seq datasets with *APOL1* genotype information.^35^ We then asked whether the mouse DEG genes were also enriched in the human RNA-seq data set. *Positive gene enrichment* refers to genes whose glomerular expression was higher in human APOL1 high-risk kidneys compared to human APOL1 low-risk kidneys. *Negative gene enrichment* refers to genes whose expression was lower in human APOL1 high-risk kidneys compared to human APOL1 low-risk kidneys.

The DEG set from HIVAN mouse model (Podo_APOL1_ HIVAN_Pos (n=330)) had positive enrichment in RNA-seq data (**Figure 4G**). In contrast, the DEG set from the mouse interferon-γ model (Podo_APOL1_ interferon-γ_Pos (n=684)) had negative enrichment in RNA-seq data (**Figure 4H**). These findings indicated that HIVAN model podocytes had similar transcriptomic signatures to the human FSGS glomeruli, reflected in human glomerular transcriptomes.

We also compared these transgenic mouse model data with urinary single-cell RNA-seq data from FSGS subjects (*ex vivo*)^37^ and with bulk RNA-seq of podocyte cell lines (*in vitro*).^38^ We conducted IPA pathway analysis based on DEGs, comparing APOL1 G1/G2 podocyte RNA with APOL1 G0/G0 podocyte RNA, using both human urine single cell data and human podocyte cell data. Both comparisons showed the EIF2 pathway as the most dysregulated pathway, with activated pathway z-score of 5.4 (urinary single cell data) (**Figure 4I**) and 2.4 (podocyte cell line data) (**Figure 4J**). Both pathway scores were similar to those of interferon-γ mouse model.

Although the interferon-γ mouse model signature in podocytes differed from that of human FSGS glomeruli, the interferon-γ mouse model may be mimicking the acute effect of APOL1 risk variants in the context of interferon-γ surge in humans, which was observed in urinary single-cell RNA-seq data and podocyte cell line data.

### Common podocyte damage patterns by APOL1-G1 in HIVAN and interferon-γ models

It is worth noting that a shared DEG set in these two models (Podo_APOL1_Pos (n=84) had a positive trend toward enrichment in human RNA-seq data, although it did not reach significance (**Figure 4K**). These genes merit further investigation, as they are potential contributors to APOL1 variant-induced damage patterns in a podocyte-specific manner and we see robust expression of *APOL1* in podocytes *in vivo* (**Figure 5A**).

**Figure 5.**
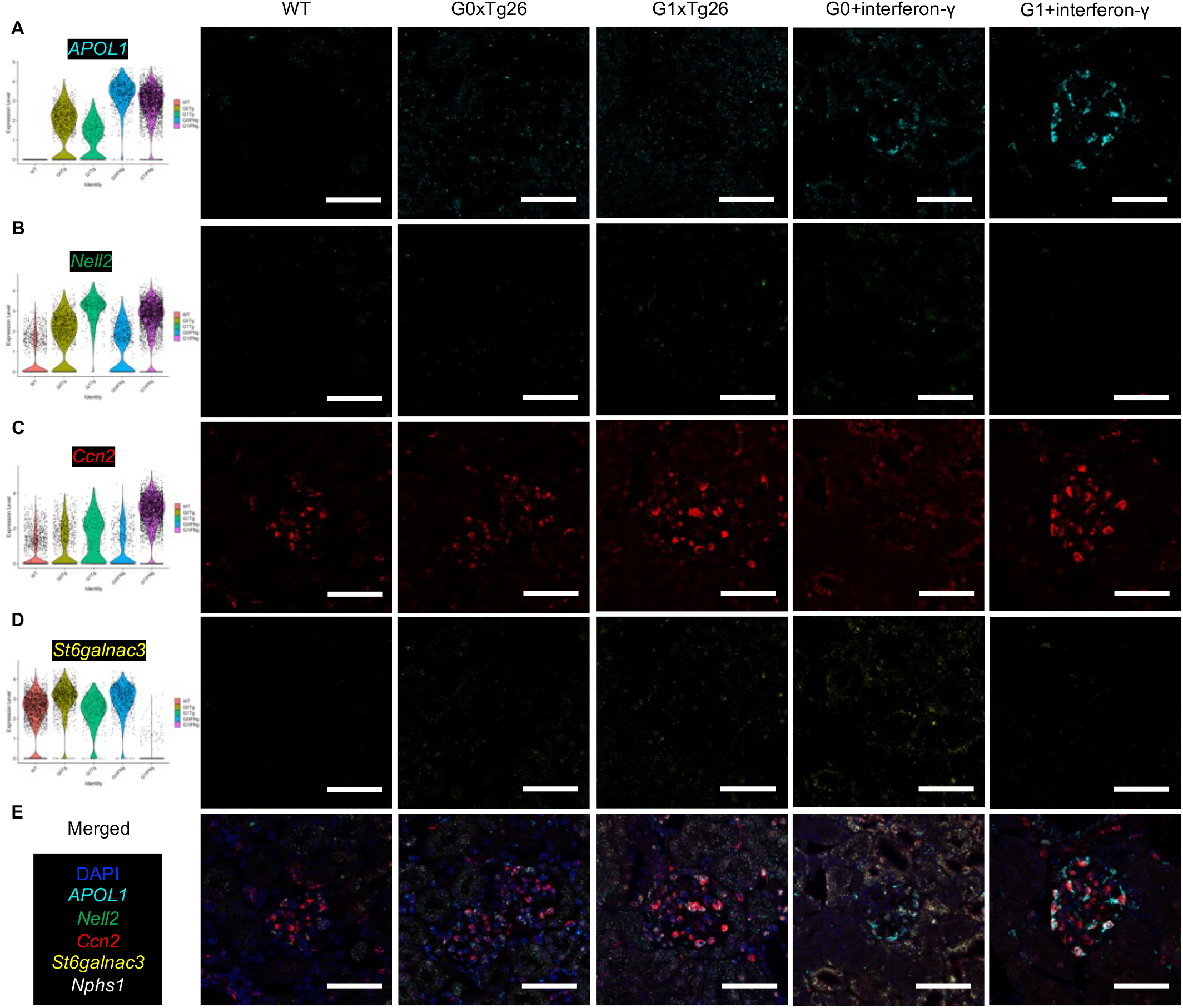
Expression of APOL1 and potential markers of podocyte injury, Nell2, Ccn2 and St6galnac3. (**A**) Violin plot showing *APOL1* expression in podocytes of snRNA-seq data and corresponding *in situ* hybridization images of *APOL1* mRNA probed in cyan color. Following interferon-γ exposure, APOL1 expression was increased in APOL-G0 and APOL1-G1 mice. (**B**) Violin plot showing *Nell2* expression in podocytes of snRNA-seq data and corresponding ISH images of *Nell2* mRNA probed in green color. (**C**) Violin plot showing *Ccn2* expression in podocytes of snRNA-seq data and corresponding ISH images of *Ccn2* mRNA probed in red color. (**D**) Violin plot showing *St6galnac3* expression in podocytes of snRNA-seq data and corresponding ISH images of *St6galnac3* mRNA probed in yellow color. (**E**) ISH overlay images showing targets including DAPI in blue color and *Nphs1* mRNA probed in white color. (Scale bars are 50 μm)

One DEG was *Nell2,* encoding neural EGF-like 1, which is an axon guidance cue; its anti-ligand is *Robo3,* expressed in endothelial cells.^40^ *Nell2* was upregulated in podocytes in both models, as shown by snRNA-seq and ISH (**Figure 5B**), indicating detachment of podocyte foot processes from GBM, as it has been shown in the similar context in neuronal synapses.^41^ Another DEG was *Ccn2* (**Figure 5C**), known pro-fibrotic factor in kidney.^42, 43^ *Ccn2* can be a marker of glomerulosclerosis. One of the DEGs downregulated in G1 was *St6galnac3* (**Figure 5D**), which is a member of the family of sialyltransferases that transfer sialic acids from cytidine monophosphate (CMP)-sialic acid to glycoproteins and glycolipids.^44^ Low *St6galnac3* expression is a marker of impaired sialylation, as shown in a human disease and a mouse model of FSGS.^45, 46^

### Cell-cell interaction analysis characterized podocyte-to-endothelial communications in APOL1 variant expressed mice

Variant *APOL1* expressed in podocytes is a major driver of podocytopathy. However, a role for other cells in glomerulopathy remains to be identified. Using single-nuclear RNA-seq data, we conducted cell-cell interaction analysis and characterized potential dysregulated interactions among cell types (**Figure 6A, 6B**). When we identified upregulated signals from podocytes to endothelial cells (**Figure 6C, 6D**), adrenomedullin (*Adm*) to calcitonin receptorlike receptor (*Calcrl*) signaling was robustly upregulated in both models, mainly by upregulation of *Adm expression* in mouse podocytes (**Figure 6E**). Adrenomedullin is a potent vasoactive substance that is produced abundantly in vascular endothelial and smooth muscle cells^47, 48^. Adrenomedullin is upregulated following podocyte injury induced by puromycin aminonucleoside^49^. Adrenomedullin has endogenous antioxidant potential to protect against ROS-induced podocyte injury.^50^ It is possible that adrenomedullin’s upregulation serves not only to protect podocytes themselves, but also to protect endothelial cells through interaction with CALCRI during APOL1-induced kidney injury. Next, we identified upregulated signals from endothelial cells to podocytes (**Figure 6F, 6G**), we found the laminin subunit alpha 3/5 *(Lama3/Lama5) – Dystroglycan 1 (Dag1)* interaction was upregulated mainly by upregulation of *Dag1* in podocytes (**Figure 6H**). *Dag1* encodes dystroglycan, which is highly abundant at the interface between the podocyte foot process and the glomerular basement membrane (GBM).^51^ Podocyte dystroglycan may be important for podocyte adhesion to GBM laminin^52^, so upregulation of *Dag1* may indicate early focal adhesion formation in APOL1-induced kidney injury.

**Figure 6.**
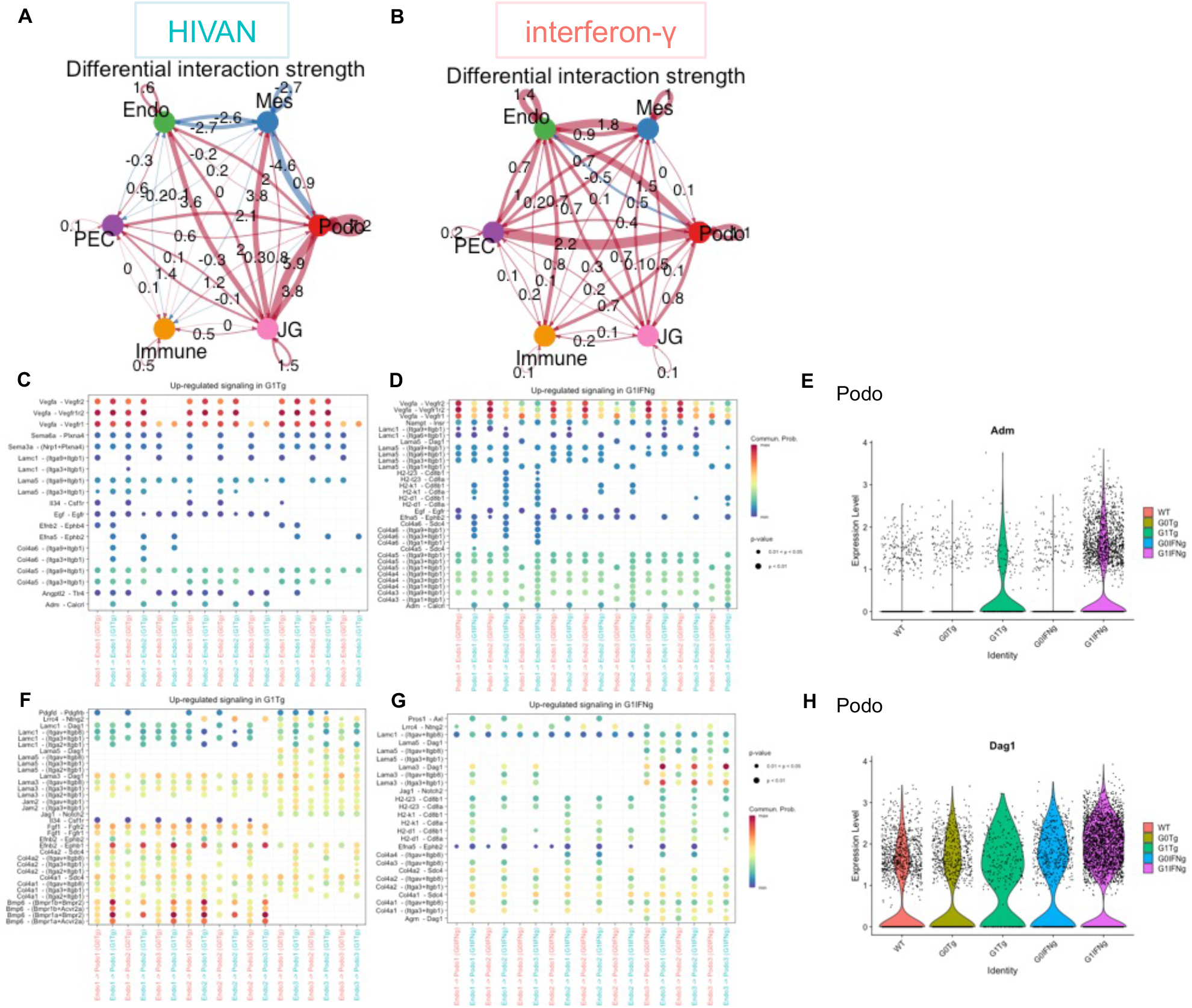
Cell-cell interaction analysis showing difference between APOL1-G1 and -G0 in two models. (**A**) Circles plot to show the differential interaction strength among cell groups between APOL1-G1 and -G0 in the HIVAN model (**B**) Circles plot to show the differential interaction strength among cell groups between APOL1-G1 and -G0 in the interferon-γ model (**C**) Dot plot to show the upregulated interactions from podocytes to endothelial cells compared APOL1-G1 and -G0 in the HIVAN model (**D**) Dot plot to show the upregulated interactions from podocytes to endothelial cells compared APOL1-G1 and -G0 in the interferon-γ model (**E**) Violin plot showing *Adm* expression in podocytes of snRNA-seq data (**F**) Dot plot to show the upregulated interactions from endothelial cells to podocytes compared APOL1-G1 and -G0 in the HIVAN model (**G**) Dot plot to show the upregulated interactions from endothelial cells to podocytes compared APOL1-G1 and -G0 in the interferon-γ model (**H**) Violin plot showing *Dag1* expression in podocytes of snRNA-seq data

### Zbtb16 can be an intermediate regulator in APOL1 variant-induced nephropathy

Transcription factors, such as WT1, can drive podocyte differentiation, and loss of certain transcription factors can be detrimental to podocyte function. We examined down-regulated DEGs in the HIVAN and interferon-γ models in both podocytes and endothelial cells (**Figure 7A**), and we identified four downregulated DEGs, *Zbtb16, Col4a3bp, Ralgapa2, Tshz2*. *Zbtb16* was recently reported to be the most important transcription factor in dexamethasone-induced podocyte protection.^53^ *Zbtb16* is induced by dexamethasone and is downregulated in collapsing FSGS. Our two mouse models showed downregulation of *Zbtb16* in both cell types, podocytes and endothelial cells (**Figure 7B-D**). This may indicate cytoskeletal disturbance and glucocorticoid-resistance in APOL1-induced nephropathy in transgenic mice, possibly in both cell types.

**Figure 7.**
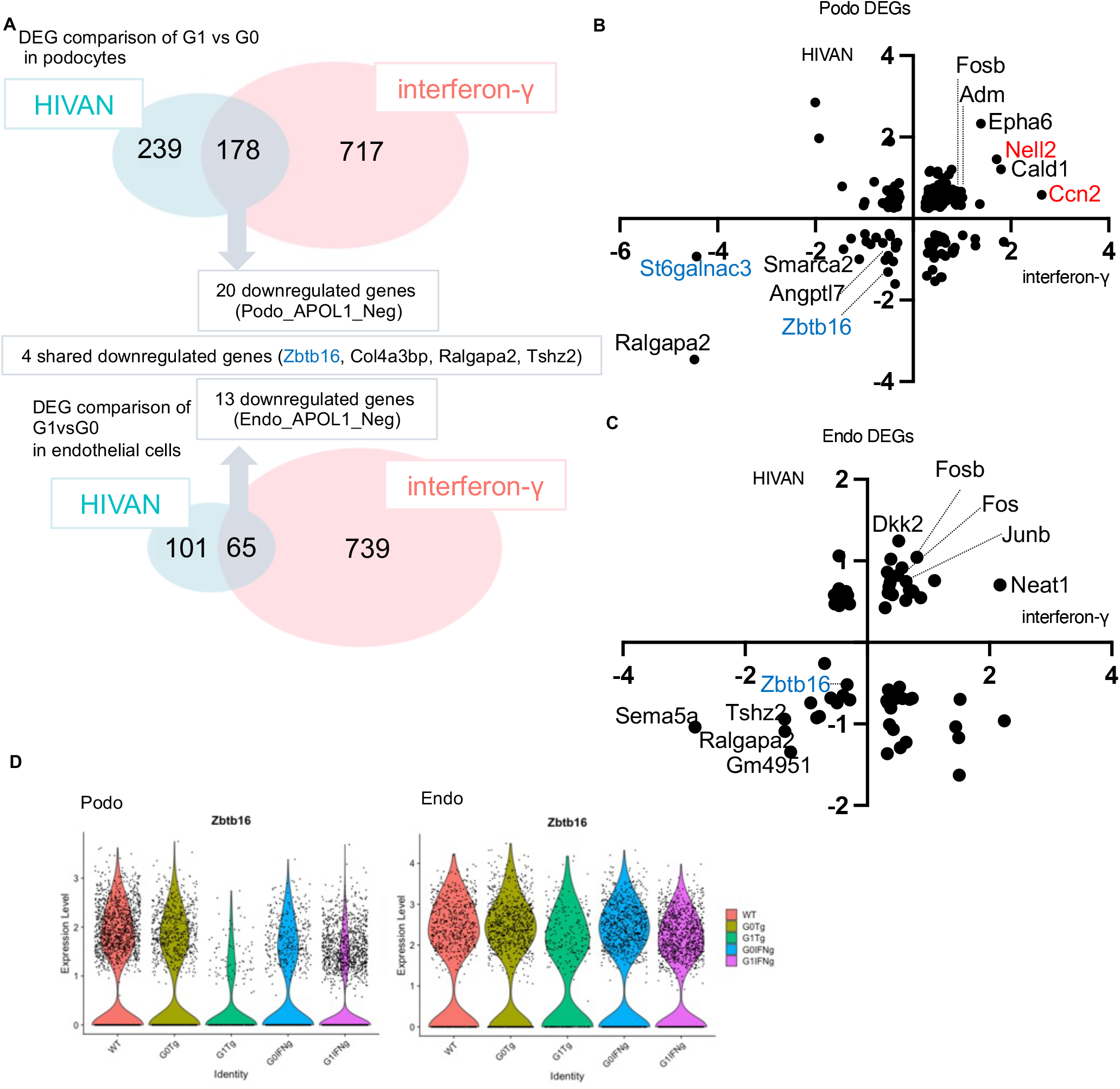
Zbtb16 was a common downregulated gene in G1 compared to G0 in two models and two cell types. (**A**) Venn diagram of DEGs, comparing APOL1 genotype (G1 vs G0) in both the interferon-γ model and the HIVAN model, in both podocytes and endothelial cells. There were four downregulated genes shared by the two models. (**B**) Scatter plot of DEGs in podocytes plotted by log2 fold change in the interferon-γ model (x-axis) and the HIVAN model (y-axis) (**C**) Scatter plot of DEGs in endothelial cells plotted by log2 fold change in the interferon-γ model (x-axis) and the HIVAN model (y-axis) (**D**) Violin plot showing *Zbtb16* expression in podocytes of snRNA-seq data (**E**) Violin plot showing *Zbtb16* expression in endothelial cells of snRNA-seq data

## Discussion

We have described transgenic mice expressing APOL1, using HIVAN mice and interferon-γ administration to BAC/APOL1 mice. Both HIV transgene and interferon-γ induced accelerated glomerular disease in APOL1-G1 compared to APOL1-G0 mice. Bulk RNA-seq of HIVAN model glomeruli showed distinct transcriptomic signatures in APOL1-G1 mice compared to both APOL1-G0 mice and HIV Tg26 transgenic mice.

APOL1-G1 and HIV gene products each caused podocyte de-differentiation. Further, single-nucleus RNA-seq of the HIVAN and interferon-γ models highlighted differences and common pathways dysregulated by APOL1-G1 compared with APOL1-G0 in these models, at cellular resolution. This unbiased approach identified DEGs which are potential disease markers or mediators of APOL1 nephropathy.

The HIVAN mouse model with human APOL1 expression from a BAC transgene provided an approach to investigate APOL1 variant-specific molecular pathways. These mice exhibited greater albuminuria induced by the combination of the HIV Tg26 transgene and APOL1-G1 gene, compared to Tg26 alone. We found greater albuminuria and fewer podocytes in APOL1-G1xTg26 mice, compared to APOL1-G2xTg26. A possible explanation for the milder phenotype in APOL1-G2 mice is lower expression of APOL1 in podocytes and glomerular endothelial cells, as shown here using multiple assays (qPCR, RNA-seq, ISH, and IHC). Clinical studies also suggest that *APOL1-G1* associates more strongly than *APOL1-G2* with HIVAN ^18^. Indeed, APO1-G1 heterozygous individuals, but not APOL1-G2 heterozygous individuals, are at increased risk for HIVAN.^54, 55^ A dual transgenic mouse model of HIVAN (the Tg26/HIVAN4 mouse), expressing APOL1-G0 or -G2 under the *Nphs1* promoter, found APOL1 expression in podocytes, but exacerbation of kidney disease compared to Tg26 mice was not observed in APOL1-G2 mice.^56^ Similarly, we found here, using a different APOL1 expression system, that APOL1-G2 did not exacerbate the HIVAN Tg26 phenotype.

The interferon-γ model was a robust model, as previously reported^16^. APOL1 mRNA and protein were upregulated in both APOL1-G0 and APOL1-G1 mice but albuminuria was induced only in APOL1-G1 mice. By comparing two different disease models, we made some novel observations concerning APOL1 nephropathy. It is well recognized that APOL1 is toxic to cells *in vitro* and *in vivo*, when expressed at supra-physiologic levels.^9, 57^ In the NEPTUNE study of patients with glomerular disease, APOL1 high-risk patients had higher APOL1 gene expression in glomeruli compared to APOL1 low risk subjects.^12^

In the present HIVAN model, we did not observe induction of APOL1 transgene expression by the HIV transgene, and HIV transgene expression was also not induced by APOL1 variants. The administration of interferon-γ to BAC/APOL1 mice represented an acute APOL1 induction model and increased APOL1 mRNA and protein expression in both podocytes and endothelial cells, to a greater extent in podocytes. Albuminuria was induced in APOL1-G1 mice but not in APOL1-G0 mice. These findings suggest that acute overexpression of APOL1-G1 in podocytes is likely inducing albuminuria.

When we compared DEGs in both models (HIVAN and interferon-γ), the enriched pathways, especially EIF2, manifested mirror images in the two models. Thus, the EIF-2 pathway was activated in the interferon-γ model, while the EIF-2 pathway was down-regulated in the HIVAN model. On the other hand, the directionality of many DEGs was also shared. Comparison with NEPTUNE glomerular RNA-seq data, gene set enrichment analysis indicated that the HIVAN model manifested a transcriptome that was close to the human (non-HIV) FSGS transcriptome, when comparing *APOL1* high-risk and low-risk genotypes. In the present report on the HIVAN mouse model, APOL1 expression induced transcriptomic changes, revealed by bulk and single-nucleus RNA-seq. HIV gene expression drives additional perturbations, such as activation of extracellular matrix related pathways. Ribosome pathway genes were downregulated by both APOL1-G1 genotype and by the Tg26 HIV transgene. The interferon-γ model is an acute APOL1 induction model. Similar human diseases have not been characterized at a molecular level. However, the model involving interferon-γ administration to BAC/APOL1mice showed EIF-2 pathway activation was similar to urine single cell RNA-seq and human podocyte cell lines, as shown above.

It may be that in human patients, *APOL1* gene induction may be either episodic or sustained leading to kidney disease. This might explain the transcriptomic differences between BAC/APOL1 mice following inferferon-γ injection model and the BAC/APOL1 model crossed with Tg26 HIVAN model.

We found several genes manifesting either increased or decreased expression in both HIVAN and interferon-γ models, which were dysregulated in APOL1-G1 mice compared with APOL1-G0 mice. If replicated in human kidney samples, these genes might serve as markers of APOL1-specific podocytopathy. For example, *Ccn2,* encoding cellular network communication factor 2, is a pro-fibrogenic factor and a possible marker to characterize podocyte fibrogenic status. *St6galnac3* is a family of sialyltransferases that transfer sialic acids to glycoproteins and glycolipids. Measuring the podocyte RNA expression of *St6galnac3* or finding defective sialylation could be an early marker to identify APOL1 variant-induced podocytopathy. How the APOL1-G1 variant induces down-regulation of *St6galnac3* requires further investigation. Similarly, by means of cell-cell interaction analysis, adrenomedullin and dystroglycan were identified as candidate markers of APOL1 variant-induced podocytopathy. These genes also warrant further investigation.

Expression of *Zbtb16* mRNA was downregulated in both podocytes and glomerular endothelial cells in both HIVAN and interferon-γ models. Glucocorticoid-resistant nephrotic syndrome has both non-genetic and genetic causes; among the latter include APOL1 high-risk variants.^58, 59^ The present finding may shed light on the mechanism of steroid-resistance in APOL1-driven kidney diseases.

We acknowledge limitations of this study. First, the BAC/APOL1 mouse has different kidney APOL1 expression levels among APOL1 genotypes, -G0, -G1 and -G2. The reasons for these variations are not known but may relate to the particular genomic location of each transgene. Differences in expression level could contribute to the APOL1-G2 mouse having a milder phenotype.

In conclusion, we have characterized APOL1 transgenic mouse models to investigate podocyte-specific genes and pathways involved in APOL1 variant-induced glomerular diseases. The HIVAN and interferon-γ models showed distinct APOL1-G1-specific transcriptomic patterns compared to APOL1-G0 mice. This identified genes differentially-expressed between APOL1-G0 and APOL1-G1 mice. These genes are potential APOL1-variant-specific therapeutic targets.

## Acknowledgements

We thank the Sequencing Facility and Bioinformatics Group (Frederick National Laboratory for Cancer Research (FNLCR), NCI, NIH) for sequencing and informatics support, Kris Ylaya (NCI/NIH) and Maria Campos (NEI/NIH) for pathological service, Patricia Zerfas (OD/NIH) for electron microscopic service, Dr. Koji Okamoto (Tohoku University, JAPAN) for scientific suggestions, Drs. Jurgen Heymann, Luis Menezes (NIDDK/NIH) for critical manuscript review. We appreciate receiving BAC/APOL1 mice from Merck Sharp & Dohme LLC, Rahway, NJ. We appreciate receiving APOL1 antibodies from Genentech, South San Francisco, CA. This work utilized the resources of the NIH HPC Biowulf cluster (http://hpc.nih.gov) and NIDDK Advanced Light Microscopy & Image Analysis Core (ALMIAC) and NIDDK Mouse Transgenic Core Facility. The content of this publication does not necessarily reflect the views or policies of the Department of Health and Human Services, nor does mention of trade names, commercial products, or organizations imply endorsement by the U.S. Government.

## Author Contributions

TY, JBK conceived the study design. TY conducted mouse experiments with support by SS. TY analyzed bulk RNA-seq data. TY conducted single-nuclear RNA-seq capture. TY and KZL analyzed single-nuclear RNA-seq data. YZ and CAW supported sequencing analysis at FNLCR/NCI. JC and SMH conducted tissue micro array creation, ISH and slide scanning. AZR assessed pathological quantification. PF assessed electron microscopic images. BS, TY, VT and PS conducted podocyte morphometry. TY drafted the manuscript and all the authors contributed for edits.

## Data Sharing Statement

Original data files and count tables have been deposited in GEO (GSEXXXX). Other data and reviewer token are available from the authors upon request.

