## Supplemental Figures for "APOL1 kidney risk variants in glomerular diseases modeled in transgenic mice"

**Supplemental Figure 1**

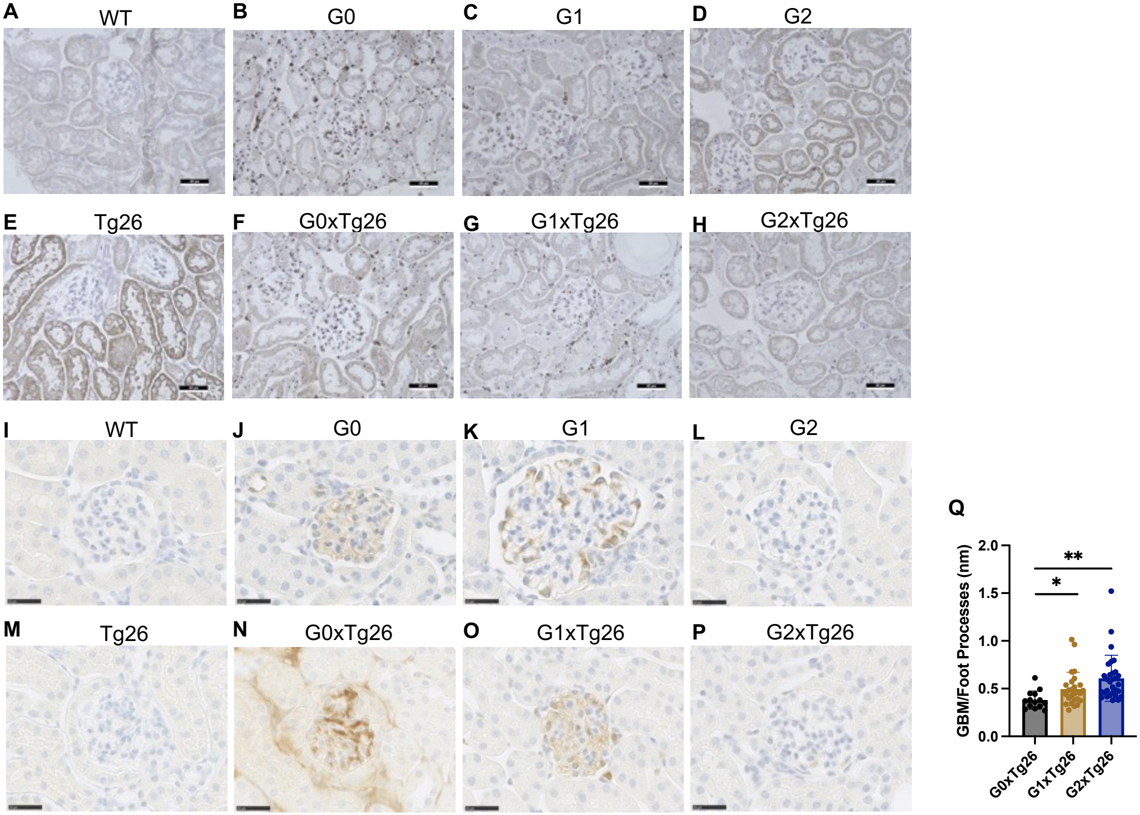


**Supplemental Figure 1. BAC/APOL1xTg26 dual transgenic HIVAN model mouse kidney**
(**A-H**) Representative images of ISH showing *APOL1* mRNA expression
(**I-P**) Representative images of IHC showing APOL1 protein expression
(**Q**) Glomerular basement membrane / foot processes (nm) measured in G0xTg26, G1xTg26 and G2xTg26 kidneys

### Supplemental Figure 2

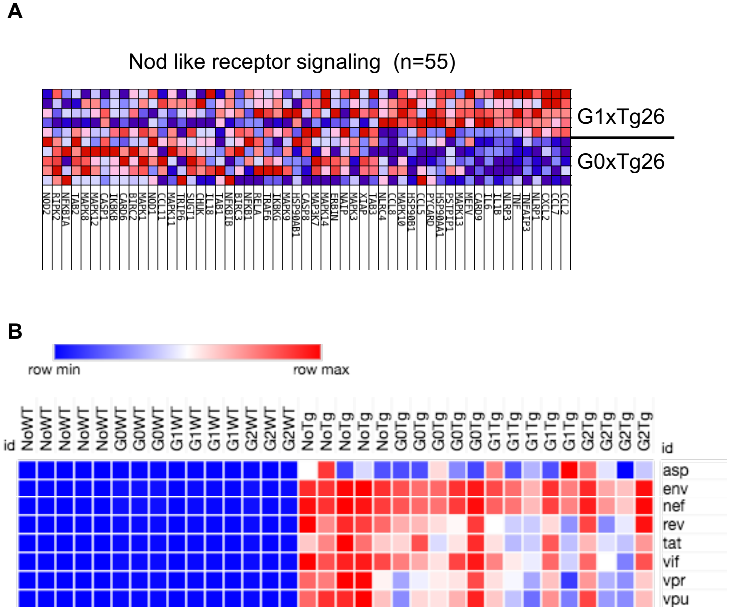


**Supplemental Figure 2. Bulk RNA-seq of BAC/APOL1xTg26 dual transgenic HIVAN mouse kidney**
(**A**) Heatmap of gene set enrichment analysis results of Nod like receptor signaling pathway comparing G1xTg26 with G0xTg26
(**B**) Heatmap of HIV gene expression of BAC/APOL1xTg26 dual transgenic HIVAN mouse kidney

**Supplemental Figure 3**

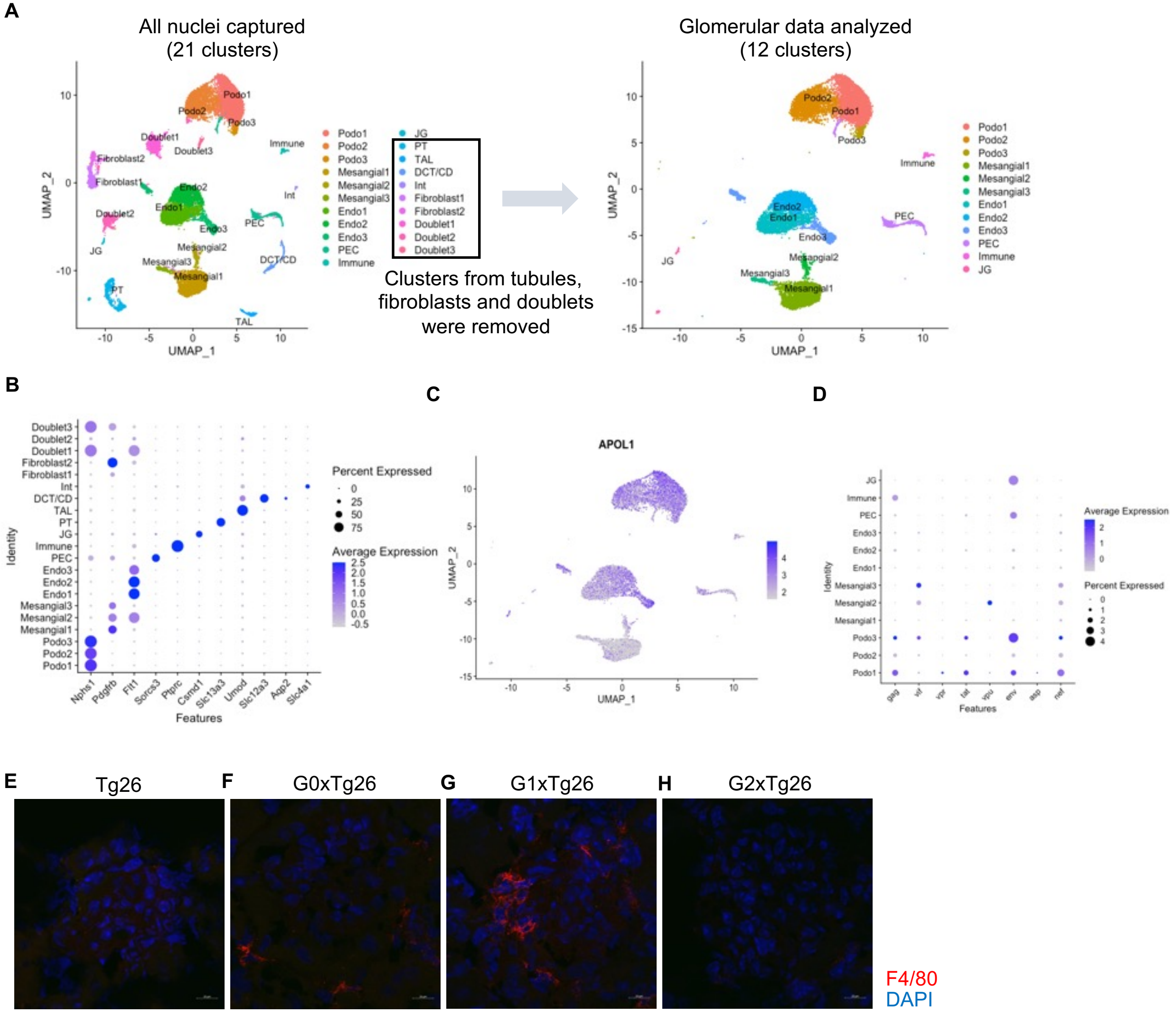


**Supplemental Figure 3. Single-nucleus RNA-seq: additional results**
(**A**) UMAP plot of single-nuclear RNA-seq data before and after the removal of clusters from tubules, fibroblasts and doublets
(**B**) Dot plot showing marker genes in each clusters before the removal of clusters
(**C**) Feature plot showing *APOL1* expression
(**D**) Dot plot showing HIV gene expression in each clusters analyzed
(**E-H**) F4/80 immunofluorescent images showing macrophage infiltration in G1xTg26 glomerulus

**Supplemental Figure 4**

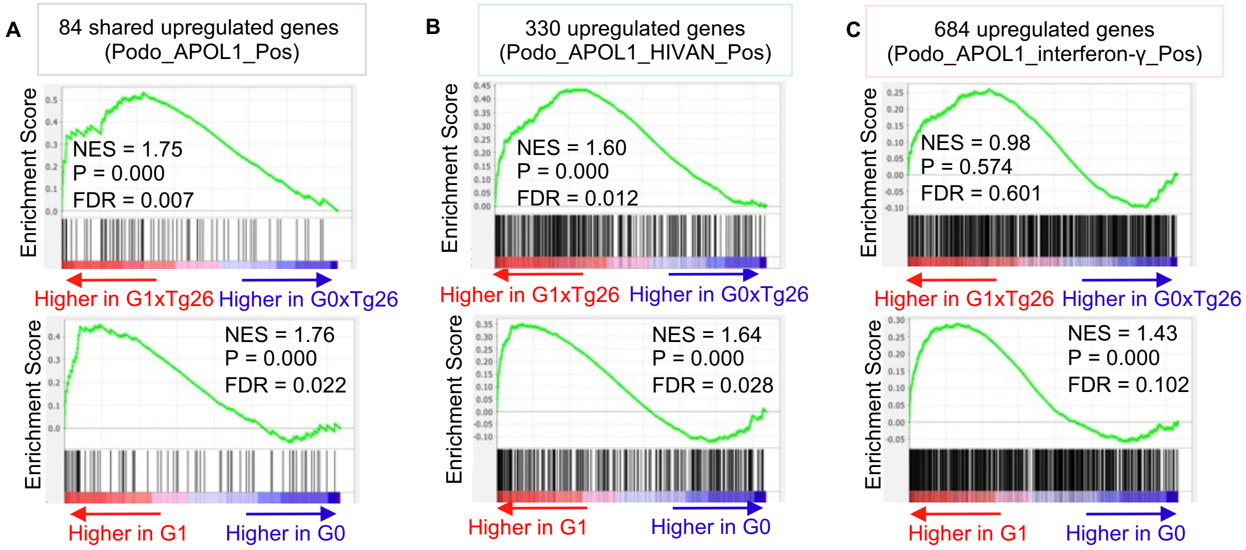


**Supplemental Figure 4. Gene set enrichment analysis results of bulk RNA-seq data using DEG sets from snRNA-seq**
(**A**) Enrichment plot of Podo_APOL1_Pos genes on bulk RNA-seq comparing G1xTg26 and G0xTg26, G1 and G2
(**B**) Enrichment plot of Podo_APOL1_HIVAN_Pos genes on bulk RNA-seq comparing G1xTg26 and G0xTg26, G1 and G2
(**C**) Enrichment plot of Podo_APOL1_interferon-g_Pos genes on bulk RNA-seq comparing G1xTg26 and G0xTg26, G1 and G2
